# ER-detect: a pipeline for robust detection of early evoked responses in BIDS-iEEG electrical stimulation data

**DOI:** 10.1101/2024.01.09.574915

**Authors:** Max A. van den Boom, Nicholas M. Gregg, Gabriela Ojeda Valencia, Brian N. Lundstrom, Kai J. Miller, Dorien van Blooijs, Geertjan J.M. Huiskamp, Frans S.S. Leijten, Gregory A. Worrell, Dora Hermes

**Author notes:** Corresponding authors: Max van den Boom, Dora Hermes.

## Abstract

Human brain connectivity can be measured in different ways. Intracranial EEG (iEEG) measurements during single pulse electrical stimulation provide a unique way to assess the spread of electrical information with millisecond precision. To provide a robust workflow to process these cortico-cortical evoked potential (CCEP) data and detect early evoked responses in a fully automated and reproducible fashion, we developed Early Response (ER)-detect. ER-detect is an open-source Python package and Docker application to preprocess BIDS structured iEEG data and detect early evoked CCEP responses. ER-detect can use three response detection methods, which were validated against 14 manually annotated CCEP datasets from two different sites by four independent raters. Results showed that ER-detect’s automated detection performed on par with the inter-rater reliability (Cohen’s Kappa of ∼0.6). Moreover, ER-detect was optimized for processing large CCEP datasets, to be used in conjunction with other connectomic investigations. ER-detect provides a highly efficient standardized workflow such that iEEG-BIDS data can be processed in a consistent manner and enhance the reproducibility of CCEP based connectivity results.

## 1. Introduction

Many studies have shown the (clinical) value of characterizing connectivity profiles during intracranial EEG monitoring with single pulse electrical stimulation (SPES). During SPES, electrode pairs are stimulated with a brief electrical pulse (1<ms) and responses are measured on all implanted electrodes. By systematic stimulation and identification of cortico-cortical evoked potentials (CCEPs), direct or indirect connectivity between electrode-sites (or structures) can be quantified. Prior studies have used CCEPs to investigate connectivity related to the motor ^1–8^, language ^2,9,18–24,10–17^, audio ^25–28^, visual ^29,30^ and limbic ^31–40^ system. CCEPs are also used in clinical settings, such as during tumor surgeries ^18,22–24,41^, vascular surgeries ^42^, and to understand and suppress seizure onset and propagation during epilepsy monitoring ^35,43–47^. This wide variety of applications, combined with release of large CCEP datasets ^48,49^, requires reliable quantification of these data to allow interpreting the CCEPs in a consistent fashion by a large audience. To provide a robust computational workflow for the processing and interpretation of CCEPs, we developed an automated framework that builds on the Brain Imaging Data Structure (BIDS; ^50,51^) and aligns with broadly established methods.

CCEPs are complex waveforms with many components. Previous studies have often taken the approach of identifying specific components from the waveform shape, which occasionally has led to different definitions of these components (reviewed in Supplementary Table 1). While some components are debated, one component – a large early response found within the first 100ms after stimulation – enjoys large consensus across studies. In recordings on the cortical surface with electrocorticography (ECoG), this response is negative, which is why it is referred to as an N1 response (illustrated in Figure 1). The N1 component is a prominently used biomarker of connectivity^1,2,17,19,21–24,26,31,42,43,3,44,45,47–49,52–56,8,57–61,9,10,13–16^, has been integrated with diffusion MRI measurement and related to the structural white matter connectome ^17,58,62^, is used to study development of this connectome ^60^, and is used for intraoperative mapping ^18,22–24^. Therefore, while CCEPs may contain more complex features, we have focused only on the detection of these early responses.

**Figure 1.**
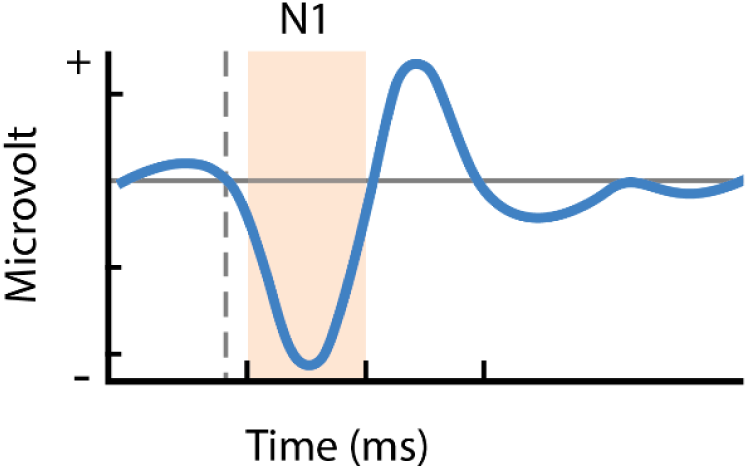
The canonical shape characterized by most studies, with the solid line indicating zero microvolt and the dashed line indicating the onset of electrical stimulation.

The identification process of the N1 response, however, is variable across studies. Clinical practice and some SPES studies rely on the manual annotation of N1s based on the visual inspection of average responses (see Table 1; ^1,3,34,43,44,46,56,63,64,13–17,19,23,24^). While some N1 responses are clearly identifiable, there is variation in the response amplitude, width and peak latency (see Supplementary Table 2 for examples). The differences in response characteristics are relevant, as they might be informative on the (in)directness and possibly the strength of the connection. However, their variation does pose a problem for the consistency and the reproducibility of results and derived conclusions. Some studies already use automated methods to detect components and thus provide some degree of standardization. However, these studies often use different methods and parameters to identify the N1 component. Table 1 illustrates how studies differ in some aspects of data processing, including baseline correction, the time window in which a deflection is considered an N1 (varying from within 5ms to 100ms) and the methods to identify N1 deflections. While all these methods are valid, the quantification of N1 responses, the scientific reproducibility of research results, and the acceptance and adaptation of evoked potential connectivity information in clinical workflows would benefit greatly from further standardization.

**Table 1.**
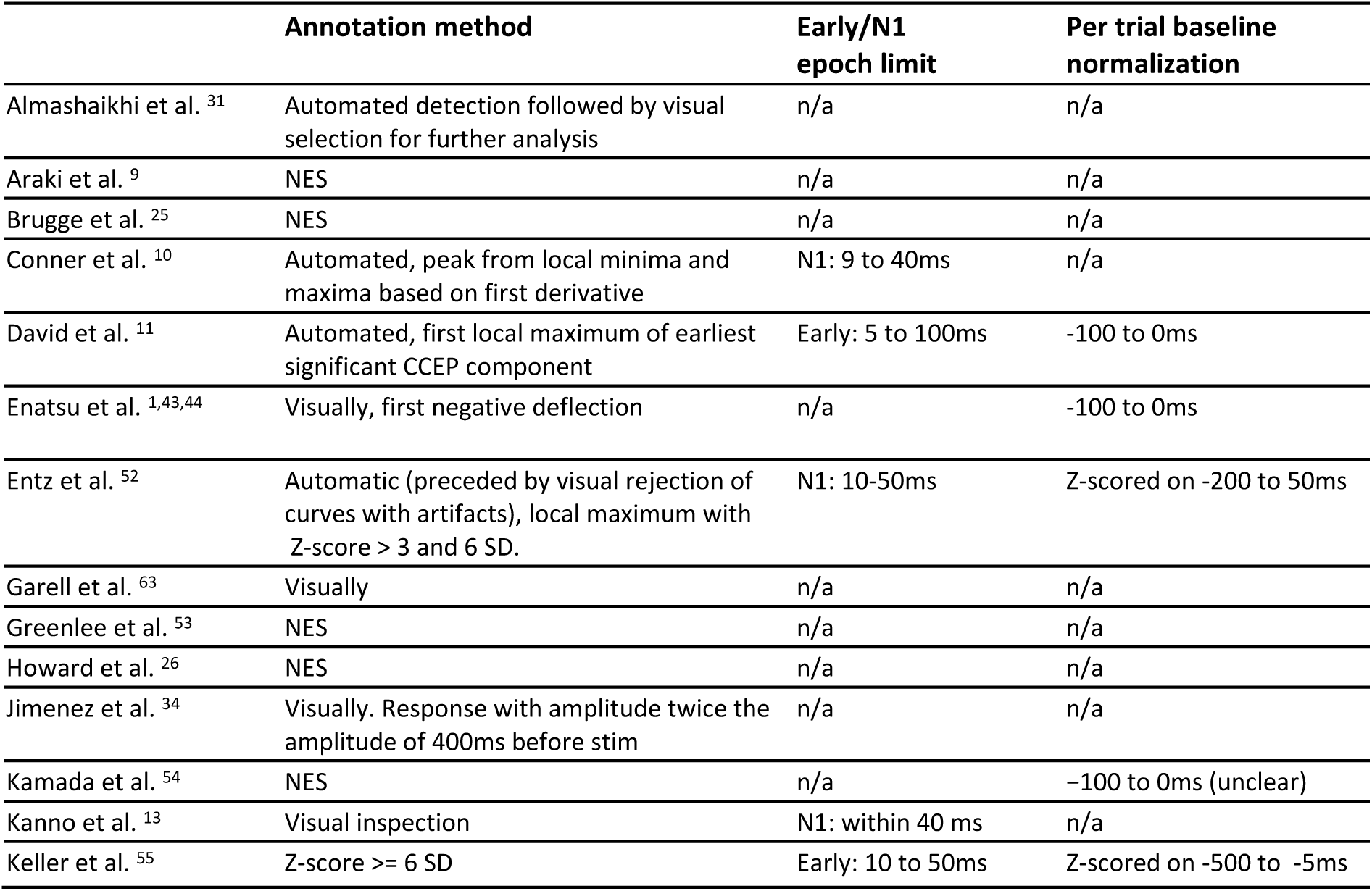
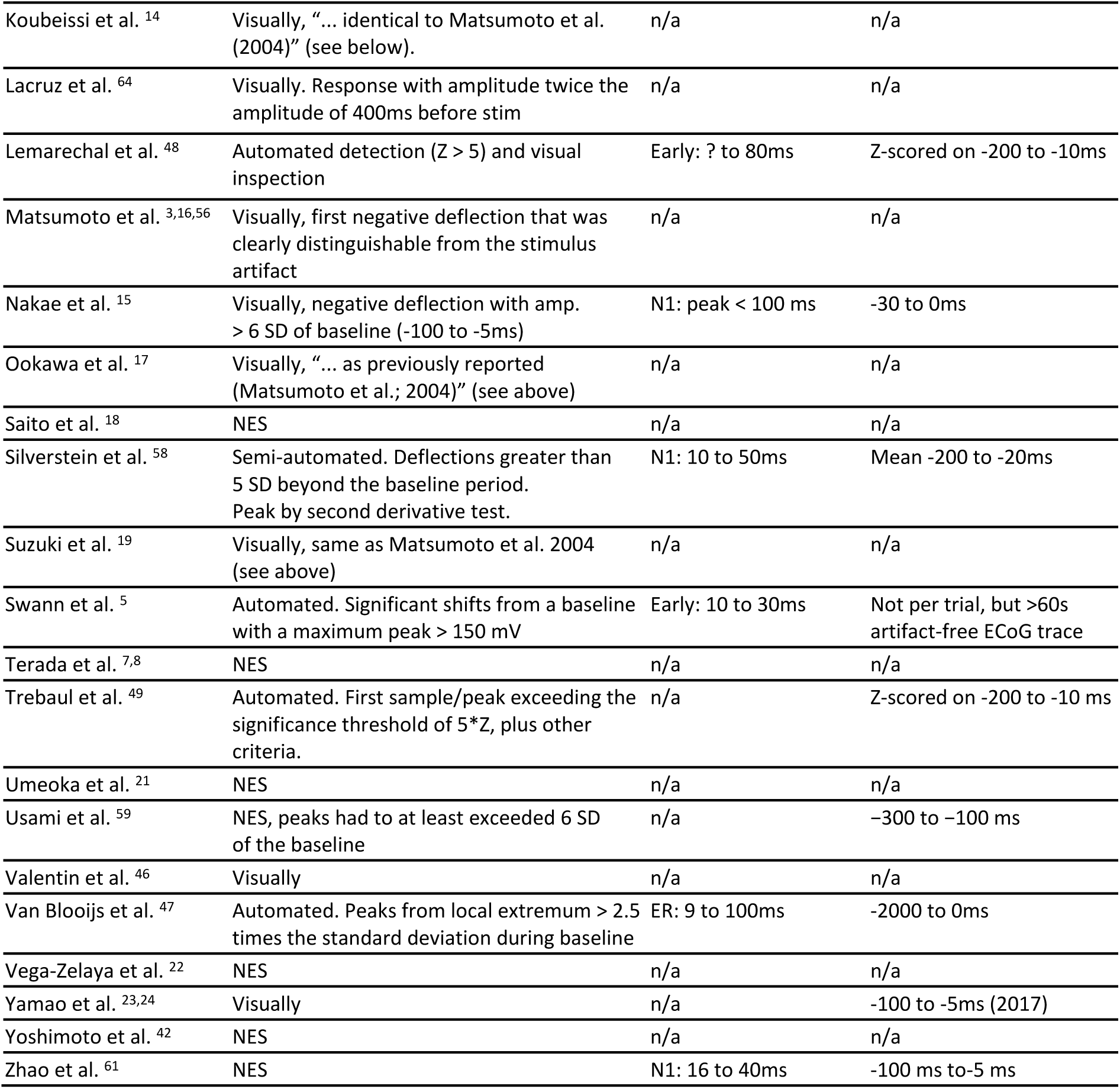
An overview of several single pulse electrical stimulation studies that use deflections in the (trial averaged) signal to identify N1/early evoked responses. If epochs are mentioned, these are relative to stimulus onset. Not Explicitly Stated (NES) indicates that the methods section did not explicitly state the method of detection (visual or computational); N/A indicates that it was either unknown or not applicable.

In an effort to standardize and facilitate fundamental and clinical studies using CCEPs, we developed a tool consisting of a Python package and BIDS Docker application that automatically identifies and characterizes these early stimulation evoked responses. It can be used to determine stimulation driven connectivity based on the N1 responses in ECoG, but also allows for the detection of positive responses that may be more common in stereo EEG (sEEG). The Early Response (ER)-detect tool processes iEEG data that is structured in the widely used BIDS standard ^50,51^, and can be run both locally by means of a graphical user interface and command-line, or in the cloud. After optional preprocessing (high pass filtering, line noise removal, re-referencing), the workflow calculates average CCEPs and detects early peak amplitudes in these responses. It can then use one of three methods to evaluate whether these early responses reach significance: ‘standard deviation from baseline’, ‘inter-trial-similarity’ or ‘wavelet filter’. Each of the three detection methods has been validated against 14 visual inspected and manually annotated CCEP datasets. After processing, ER-detect provides standardized output on the evoked responses, including a data file for further processing, visualizations of the converging (incoming) responses per electrode, visualizations of the divergent (outgoing) responses per stimulated-pair, and visual representations of the response latency and amplitude in connectivity matrices.

## 2. Results

### 2.1 Early response detection using the application

ER-detect is open-source and freely available as a Python package (with optional GUI) and Docker image, which can run locally on any operating system (e.g. Windows, Mac or Linux) or cloud environment. Paragraph 2.1.1 below describes how the CCEP input data is provided and paragraph 2.1.2 shows the output after preprocessing and evoked response detection. The tool generates a data file for further processing and visualizations that can be used for interpretation.

#### 2.1.1 Input

The tool expects the input to be formatted according to BIDS standard (https://bids-specification.readthedocs.io/ ^50,51^). An example of an iEEG-BIDS data structure with a minimum amount of files is provided in Figure 2.

**Figure 2.**
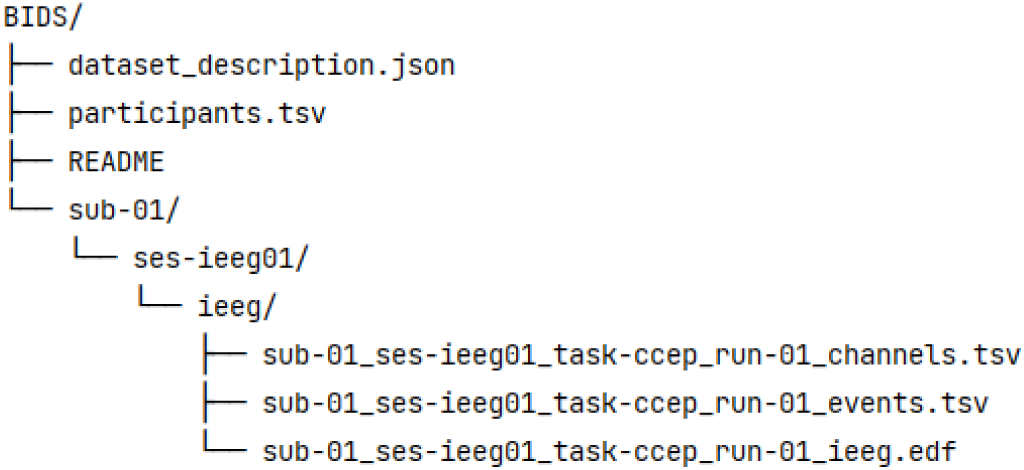
An example of a minimal BIDS structure with one participant (sub-01) and one iEEG dataset.

Multiple participants and runs that are in the BIDS root folder can be processed consecutively and can be selected using the GUI (see 2.1.3) or by passing command-line arguments. In accordance with the BIDS specification, the iEEG signal data can be stored in several widely used formats: European Data Format (EDF), BrainVision or MEF3 (.mefd). The ‘*_channels.tsv*’ should contain the iEEG channel details as required by the iEEG-BIDS specification and the ‘*_events_tsv*’ should contain the stimulation details as suggested in the specification ^50^.

#### 2.1.2 Output

The tool produces two types of output: a data file for further analyses, and visualizations for quick interpretation. The data file stores the configuration that was used, the electrode-pairs that were stimulated, the electrodes that were measured, the average signal over stimulation trials, and - importantly - the early response detection results. These results contain a connectivity matrix that indicates, for each stimulated electrode-pair and each measured electrode, whether an early response is detected. When indeed such a deflection is found, the peak latency (in milliseconds from stimulus onset) and peak amplitude (in µV) of the deflection are stored as well.

The visual output offers several views that depict: the incoming connections, the outgoing connections, and the connectivity matrix with amplitude and latency. The incoming connections are represented in a set of images with one image per measured electrode. Each image shows the average signal and evoked responses for one measured electrode, given each stimulated electrode-pair (see Figure 3A). The outgoing connections are represented in a set of images with one image per stimulated electrode-pair. Each image shows, for one stimulated electrode-pair, the average signal and evoked responses on each of the measured electrodes (see Figure 3B). The connectivity matrices provide a single overview of all the stimulated and measured electrodes, one matrix depicts the peak amplitudes of the evoked deflections (see Figure 3C), and one depicts the latency of the evoked deflection peak (see Figure 3D). The connectivity matrices provide a quick overview of the connectivity in the brain based on CCEPs.

**Figure 3.**
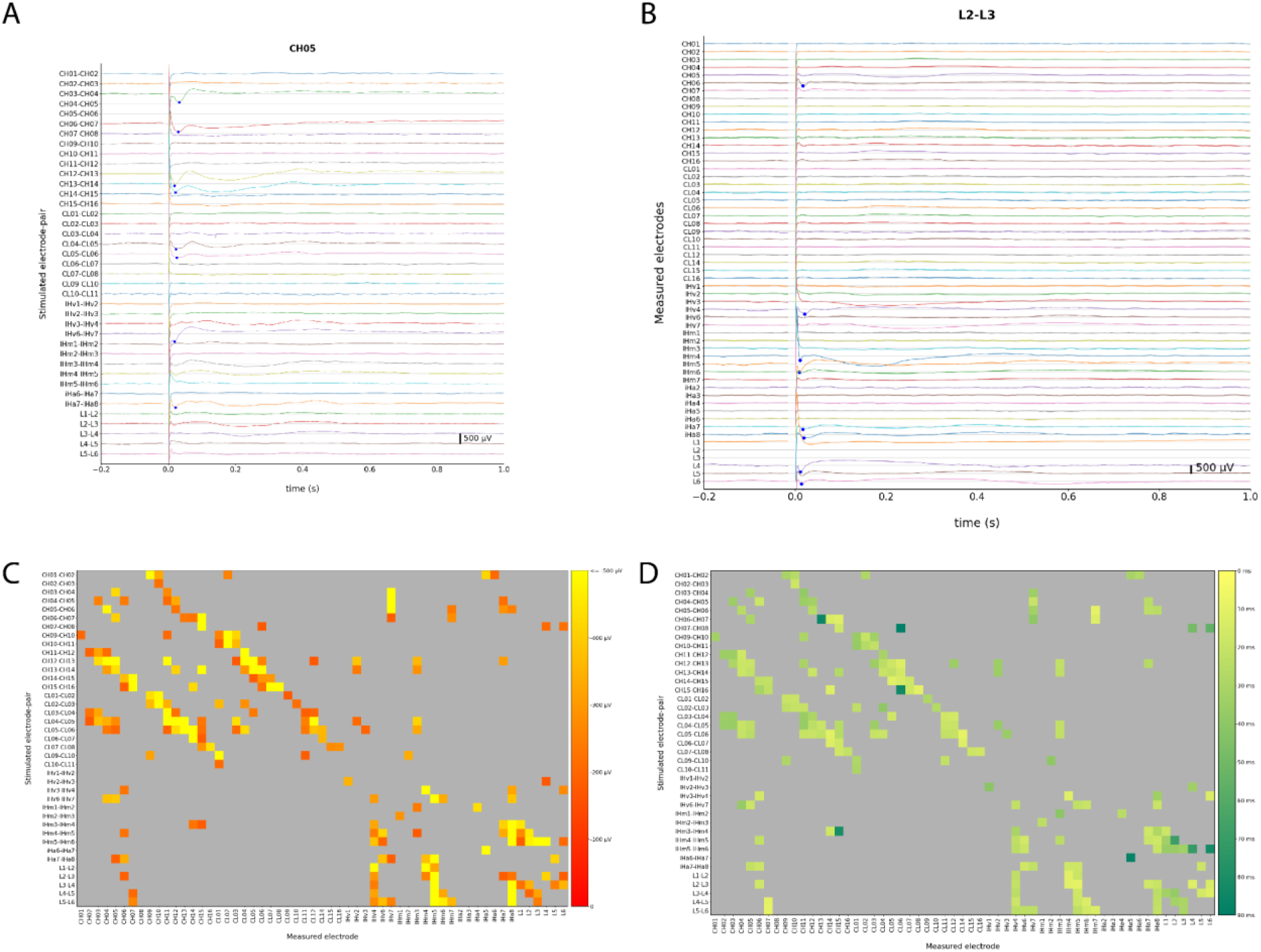
Cortico-cortical evoked potential visualizations of subject ‘UMCU21’. (A) The incoming connections for one of the measured electrodes. The blue dots mark the peaks of the negative deflections that were detected. (B) The outgoing connections for one stimulated electrode-pair, part of a set of images. (C, D) Connectivity matrices here gray cells indicate no evoked deflection and colored cells indicate (C) the peak amplitude of the deflection, or (D) the latency of the deflection peak.

These outputs can be obtained with a number of configuration settings in the tool. Several preprocessing steps, such as high-pass filtering, re-referencing and line-noise removal, can be applied to the CCEP signal data by configuration in the GUI or by passing combinations of command-line arguments. After preprocessing, early responses are detected. The outputs in Figure 3 were obtained using the ‘deviation from baseline’ method, which is similar to what many other studies used before ^5,15,60,64,34,47–49,52,55,58,59^, by identifying a deflection as an evoked response only when (the peak of) a local extremum passes a certain threshold. The ‘deviation from baseline’ method uses a default threshold of 3.4 times the standard deviation during baseline (default from -1000 to -100ms). Two other detection methods are available based on inter-trial similarity and wavelet filtering. All preprocessing, detection and output parameters can also be defined in a JSON configuration file according to the clinical, or scientific context. A complete list of configuration options is available at: https://github.com/MultimodalNeuroimagingLab/erdetect/wiki/Configuration.

#### 2.1.3 Graphical User Interface

The python package offers a convenient graphical user interface that allows the user to select a BIDS input folder, easily filter and select the subject and iEEG datasets that require processing, configure pre-processing, detection, epoching, and visualization settings (Figure 4).

**Figure 4.**
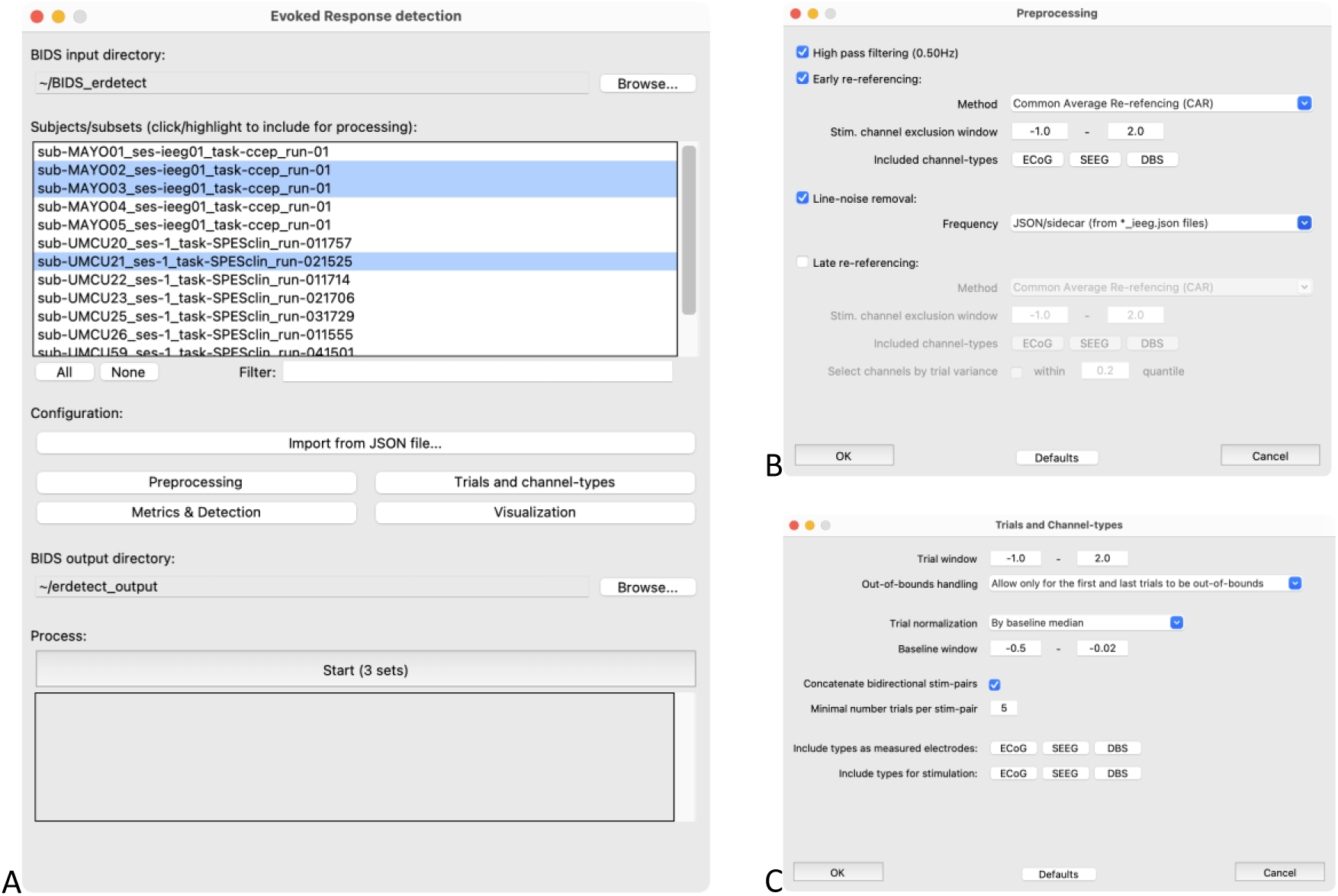
A graphical user interface is available in Windows, MacOS and Linux. Panel A shows the main window at which the input/output folder, subjects and datasets for processing and detection can be selected. The preprocessing options (panel B), trial and channel options (panel C), detection settings and visualization settings can all be modified.

#### 2.1.4 Optimized performance for speed and memory

Modern iEEG datasets are often recorded at a high sampling rate (e.g. at least at 1kHz) and include a large number of recording sites (e.g. 128 channels). While such rich datasets can be scientifically advantageous, processing them can prove challenging, either by limits in computational power or memory. The library employs routines that, depending on the (pre-)processing options, are optimized for processing speed and memory usage (results are shared at https://github.com/MultimodalNeuroimagingLab/ieegprep/wiki).

### 2.2 Validation of the early response detection

To validate the early response detections from the app, we compared the automated detected responses to the manual annotation results of each of the 14 visually inspected datasets. However, before comparing the automatic detection to the visual/manual annotations, we first investigated the inter-rater reliability (paragraph 2.2.1). The consensus between manual annotations provides a ceiling for a comparison between the automatic detection and manual annotations (paragraph 2.2.2).

#### 2.2.1 Inter-rater reliability of manual annotations

Different raters from two hospitals (see Table 4) annotated the ECoG datasets and indicated whether a negative early response (i.e. N1) was present in the average evoked response. The inter-rater reliability shows that there is a strong consensus between raters on the responses considered as not having an N1 deflection (specificity: 96%; Table 2). However, as expected, there is less agreement on which responses should be considered as having an evoked N1 deflection (sensitivity 68%; Table 2). It is important to note that over all datasets there are significantly more average traces that were considered to have no evoked responses (53 223) than as having an evoked responses (5 955). As such the precision was also calculated and found to be at 68%. The average Cohen’s kappa value over all annotations was .60.

**Table 2.** Annotation descriptives and inter-rater reliability of evoked responses. Total evoked responses (ER) and total non-ER annotations were reported for each rater. Measurements include accuracy ((TP+TN)/total), sensitivity, specificity and Cohen’s Kappa.

| Subject | Rater 1 | ER |  | Rater 2 | ER |  | Accuracy | Sensitivity |  | Specificity |  | Kappa |
| --- | --- | --- | --- | --- | --- | --- | --- | --- | --- | --- | --- | --- |
|  |  |  | non-ER |  |  | non-ER |  | 1vs2 | 2vs1 | 1vs2 | 2vs1 |  |
| MAYO04 | MvdB | 18 (7%) | 226 (93%) | SB | 19 (8%) | 226 (92%) | 96% | 72% | 72% | 98% | 98% | .70 |
| MAYO05 | MvdB | 44 (1%) | 3652 (99%) | SB | 41 (1%) | 3700 (99%) | 99% | 73% | 78% | 100% | 100% | .75 |
| UMCU20 | DvB | 90 (13%) | 607 (87%) | MvdB | 66 (9%) | 631 (91%) | 92% | 56% | 76% | 97% | 94% | .60 |
| UMCU21 | DvB | 198 (13%) | 1274 (87%) | MvdB | 85 (6%)<sup>1</sup> | 1432 (94%)<sup>2</sup> | 90% | 35% | 82% | 99% | 91% | .44 |
|  |  |  |  | SB | 62 (4%) | 1386 (96%) | 89% | 27% | 85% | 99% | 89% | .36 |
|  | MvdB | -<sup>1</sup> | -<sup>2</sup> |  |  |  | 97% | 62% | 85% | 99% | 98% | .71 |
| UMCU22 | DvB | 1032 (21%) | 3772 (79%) | MvdB | 494 (10%) | 4300 (90%) | 86% | 41% | 85% | 98% | 86% | .48 |
| UMCU23 | DvB | 385 (18%) | 1711 (82%) | MvdB | 412 (20%) | 1681 (80%) | 88% | 71% | 67% | 92% | 93% | .62 |
| UMCU25 | DvB | 220 (8%) | 2515 (92%) | MvdB | 260 (10%) | 2472 (90%) | 93% | 65% | 55% | 95% | 97% | .55 |
| UMCU26 | DvB | 688 (30%) | 1604 (70%) | MvdB | 314 (14%) | 1973 (86%) | 82% | 43% | 95% | 99% | 80% | .50 |
| UMCU59 | JvdA | 103 (6%) | 1584 (94%) | MvdB | 100 (6%) | 1587 (94%) | 98% | 85% | 88% | 99% | 99% | .86 |
| UMCU62 | JvdA | 368 (9%) | 3898 (91%) | MvdB | 257 (6%) | 4006 (94%) | 96% | 59% | 85% | 99% | 96% | .68 |
| UMCU67 | JvdA | 485 (10%) | 4378 (90%) | MvdB | 214 (4%) | 4649 (96%) | 93% | 39% | 87% | 99% | 94% | .50 |
| **Total** |  | → | → |  | **5955** | **53223** |  |  |  |  |  |  |
| **Average** |  | → | → |  | **(10%)** | **(90%)** | **92%** |  | **68%** |  | **96%** | **.60** |

#### 2.2.2 Validation of automated detection methods

All three automated detection methods were validated against the visual inspected and manually annotated data. Figure 5 illustrates the performance of each method on a group level, Supplementary Figure 1 provides the ROC curves per participant.

**Figure 5.**
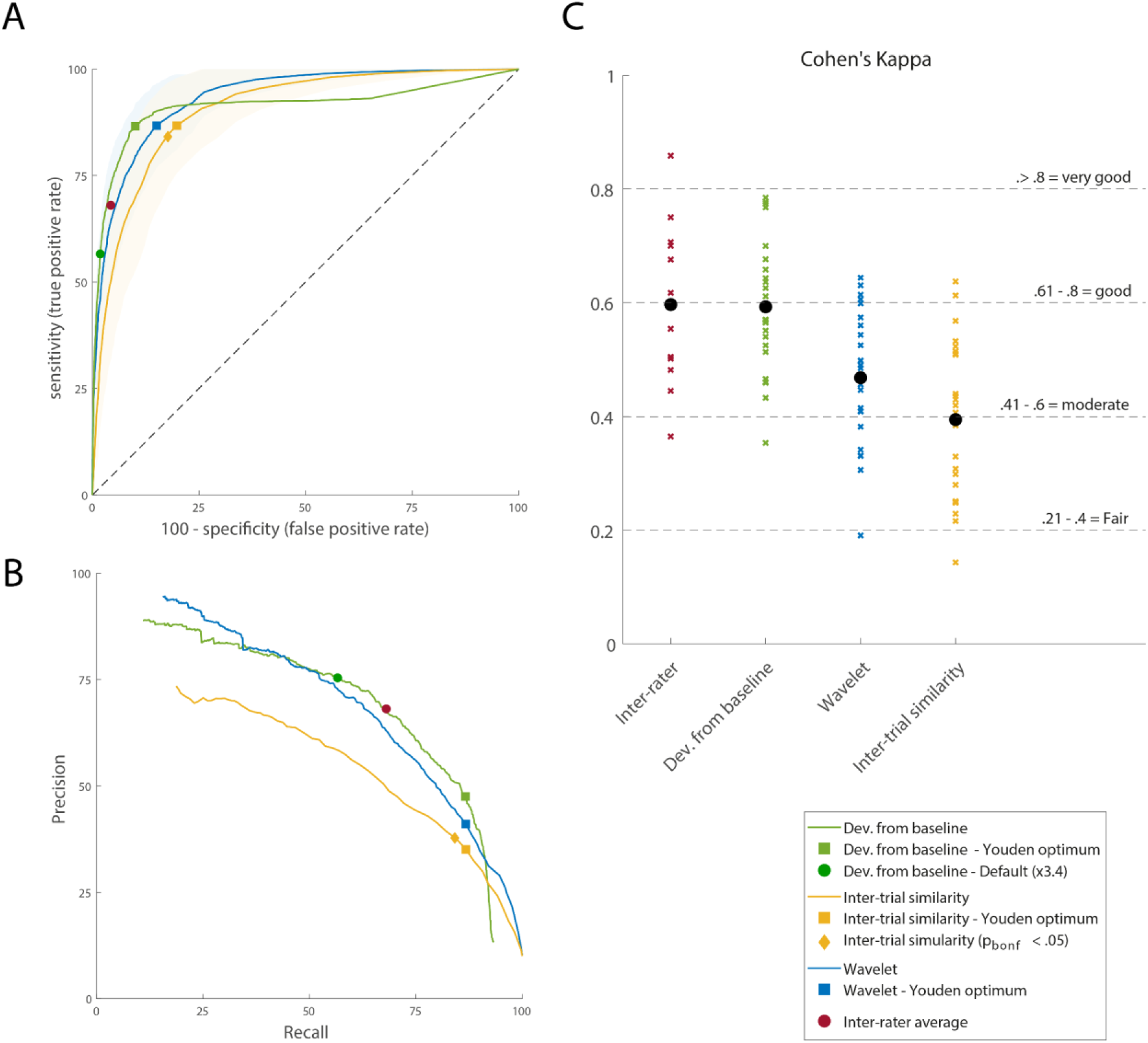
Performance of the three different detection methods**. A)** ROC curves projecting the true positive rate (sensitivity) and false positive rate (100-specificity) for each automated detection method. The ‘deviation from baseline’ method is shown in green with the threshold (as a factor of the standard deviation during baseline) ranging from 1 to 1500. The green dot represents this method’s performance based on the default setting of 3.4. The ‘inter-trial reliability’ performance is shown in yellow with threshold t-values ranging from -30 to 30. The ‘wavelet’ performance is shown in blue with power thresholds ranging from 0 to 60 000. **B)** Precision-recall curves project the Positive predictive value against the recall (sensitivity). The same ranges are applied as in panel A. **C)** The Cohen’s Kappa values of the inter-rated annotations and their comparisons on each of the three detection methods. Each annotation is shown as a cross with the average over annotations as a black dot.

The default detection method, which is set to evaluate a peak at more than 3.4 times the standard deviation from baseline as an evoked response, resulted in an average sensitivity of 57%, a specificity of 98% and a kappa of .59. The Youden optimum (at 1.7 times the standard deviation from baseline) was more sensitive, but less specific and therefore may be used if a less conservative threshold is preferred for the detection of early responses. The ‘wavelet filter’ method, based on a Youden optimum (cutoff power^2^ = 1100), yielded a mean sensitivity of 87%, a specificity of 85% and a kappa of .47. The ‘inter-trial similarity’ method, based on its Youden optimum (cutoff t = 4), resulted in a mean sensitivity of 87%, a specificity of 80% and a kappa of .39.

The results on the ’deviation from baseline’ method are similar to what was found during manual annotation (see 2.2.1 above), having almost the same Cohen’s kappa value, a very high specificity, and a medium sensitivity. These results imply - in the same way as between raters - that when there is no evoked response, both the rater and the algorithm agree. However, when there is an evoked response, the automated detection might not agree exactly with a rater.

### 2.3 Example with sEEG data

Recent iEEG practices show a gradual movement from ECoG to sEEG recordings for the localization of epilepsy sources ^65–67^. We therefore tested how the early response detection performs on sEEG data. In sEEG data, the early responses can be positive or negative, depending on the location of the electrode compared to the cortical surface. Figure 6 shows how, in an openly available sEEG dataset with single pulse stimulation (data available on: https://doi.org/10.18112/openneuro.ds004457.v1.0.2; ^29^), the ER-detect tool can detect both positive and negative responses while applying adjusted common average re-referencing ^68^. Note that both negative and positive detections are stored in the output data file for further post-processing. For example, in Figure 6, the stimulation electrodes RB1-RB2 induced both a negative and a positive response in the measured electrode RB4. In these cases, the user can further choose how to interpret each of these early responses.

**Figure 6.**
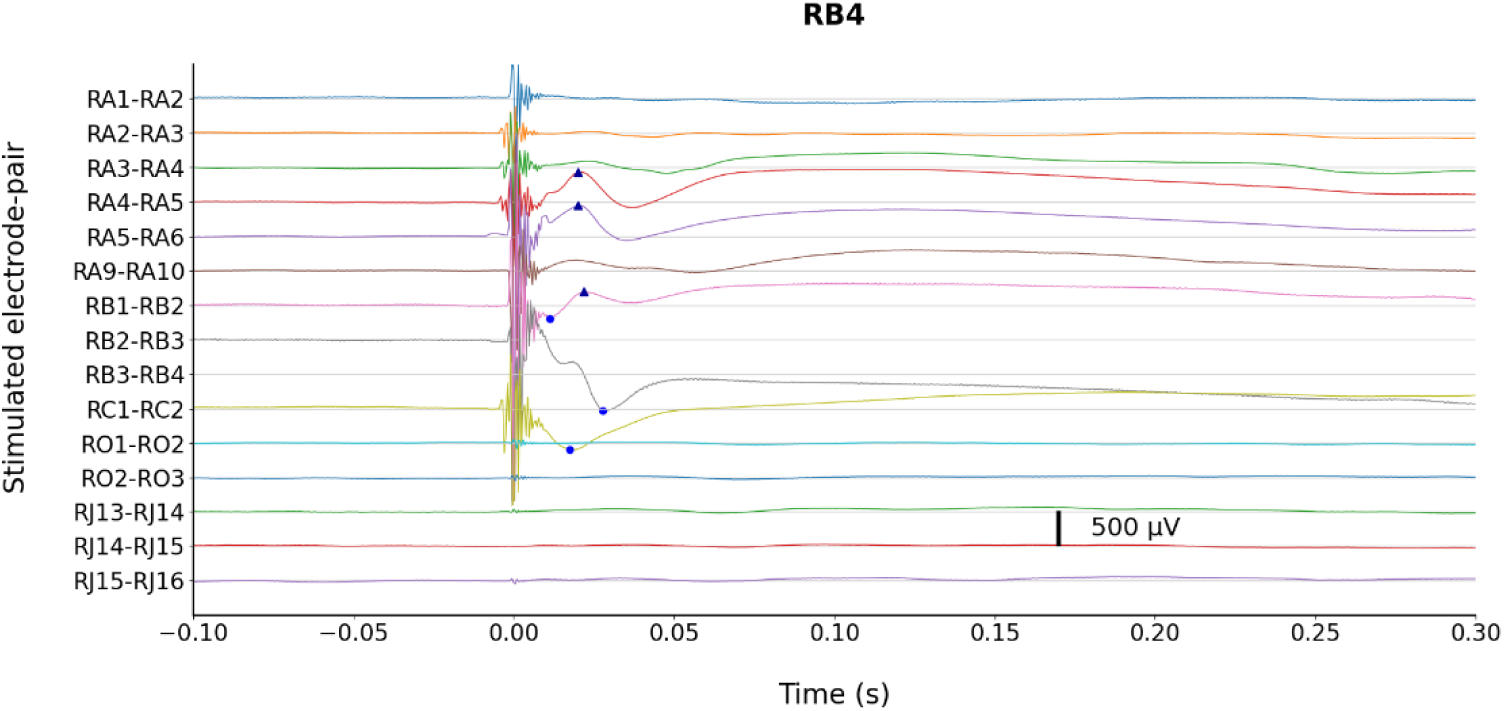
Visualization of the detection results for both positive and negative deflections in sEEG data (‘sub-05’, public dataset from Huang et al, 2023). The lighter blue dots mark the peaks of the negative deflections that were detected, while the darker blue triangles signify the peaks of the positive deflections. This figure shows an exemplary subset of the results, but other stimulation and measurement are available online for further investigation. Adjusted common average re-referencing was applied by ER-Detect based on 20% of the channels with the lowest trial signal variance (i.e. the ‘--late_reref_CAR_by_variance’ flag).

## 3. Discussion

The ER-detect tool presented in this article provides a robust workflow to preprocess BIDS structured iEEG data collected during single pulse electrical stimulation and detect early evoked responses in a fully automated and reproducible fashion. The detected responses are visualized for the immediate interpretation of cortical connectivity and are stored in connectivity matrices for further analyses. With the use of this application, iEEG data can be processed in a more consistent way, improving the reproducibility of CCEP connectivity results. All methods of detection were validated against visually inspected and manually annotated data from two different sites by four independent raters. Automatic detection by the tool proved to be similarly reliable as manual annotations between these raters.

The reliability of identifying negative evoked responses (i.e. N1s) by visual inspection was established on 14 iEEG datasets from two hospitals. The inter-rater reliability yielded a moderate Cohen’s kappa (.60). While agreement on the signal without an evoked response was high and evident (specificity: 96%), consensus on the signal with an evoked response was more problematic (sensitivity: 68%). The relatively high disagreement amongst raters, on what is considered a typical evoked response, confirms the problem found in practice. Namely, the subjectivity and the variety of ways that evoked responses are identified by different research groups, or by individuals within the same group. Not all annotators in our study had extensive clinical expertise with CCEP data.

However, a previous study found that non-experts had similar inter-rater agreement compared to experts in the setting of fiber tract segmentation ^69^. It is thus unlikely that the inter-rater scores could be further improved by employing experts or further training. This confirms the opportunity to reduce rater inconsistencies caused by human bias and improve the reproducibility by standardization and automated detection.

The automatic detection with the ER-detect tool performs on par with human annotation. In particular, the default method, that relies on peak deviation from baseline, performed very close to human identification of negative evoked deflections with a moderate Cohen’s kappa (.59), high specificity (98%) and medium sensitivity (57%). This method of baseline deflection aligns well with the implementations of automatic detection in many other studies (see Table 1; ^5,15,60,64,34,47–49,52,55,58,59^). The advantage of the ER-detect tool is that the preprocessing and parameters are standardized, detection can be run in a platform independent manner and is therefore fully reproducible.

Besides an N1 response, CCEPs can also contain more complex responses. The ‘deviation from baseline’ method assumes that the CCEP waveform has an N1 component. Other methods without such assumptions are available to analyze CCEPs, some of which are not yet implemented in ER-detect. These include methods based on the reliability of responses ^29,38,70,71^, or spectral analyses to explore stimulation induced power changes between the 70 to 200Hz range ^20,54,59,72^. ER-detect can be used in conjunction with such other methods and potentially allow quantitative comparisons to better understand the different features of human brain connectivity. Yet, if a user is more interested in reliable responses, the ‘inter-trial similarity’ method in the tool can provide insight into the CCEP waveform without presumption of the N1.

The ER-detect tool can provide a robust solution for preprocessing iEEG data that are structured according to BIDS and the detection of early responses in the CCEPs. It has been optimized for usability in several ways, including computational efficiency, a GUI, and implementation as a BIDS-app. As such, it provides an accessible and *easy-to-use* solution for a broad audience, already allowing others to freely explore openly available CCEP datasets ^29,38,48,49,60^ in a reproducible manner. This reproducible workflow allows the connectivity measures from these unique iEEG data to be further integrated with other large scale connectivity studies.

## 4. Methods

### 4.1 Participants

We included data from 14 participants who underwent clinical ECoG monitoring preceding epilepsy surgery and SPES for clinical decision making from two different sites. Five participants were patients at Mayo Clinic (Rochester, Minnesota, USA), where the study was approved by the Institutional Review Board (IRB) of Mayo Clinic (IRB #15-006530) and patients provided informed consent to participate in the study, in accordance with the declaration of Helsinki (2013). Nine participants from the University Medical Center Utrecht (UMC; Utrecht, The Netherlands; METC18-109) from a previously published dataset ^60^ were included in the study. In this previously published dataset, subjects who underwent epilepsy surgery in the UMC Utrecht between 2008 and 2020 were included in a retrospective epilepsy surgery database ^73^, with approval of the Medical Research Ethical Committee of UMC Utrecht. For subjects included between January 2008 and December 2017, the Medical Research Ethical Committee waived the need for informed consent. Since January 2018, subjects were explicitly asked for informed consent to collect their data for research purposes. Table 3 provides an overview with the patient demographics.

**Table 3.** Patient details and electrode coverage. In the number (#) of electrodes columns, total indicates the number of electrodes that were placed. Stim and Rec indicate the number of stimulated electrode-pairs and recording electrodes that were included for analysis. Location reports on the hemisphere (Left and/or Right) and the lobes that are (partially) covered by electrodes.

| Patient | Age | Sex | # ECoG electrodes |  |  | # sEEG/Depth electrodes |  |  | Location |
| --- | --- | --- | --- | --- | --- | --- | --- | --- | --- |
|  |  |  | Stim | Rec | Total | Stim | Rec | Total |  |
| MAYO01 | 16 | F | 38 | 64 | 76 | 0 | 12 | 12 | Left parietal, occipital, temporal |
| MAYO02 | 41 | M | 47 | 56 | 56 | 3 | 8 | 8 | Left frontal, parietal, temporal |
| MAYO03 | 62 | M | 26 | 34 | 36 | - | - | 0 | Right frontal, parietal |
| MAYO04 | 65 | F | 16 | 21 | 24 | - | - | 0 | Left frontal, parietal |
| MAYO05 | 36 | M | 54 | 83 | 84 | 0 | 12 | 12 | Left frontal, parietal, temporal |
| UMCU20 | 11 | M | 9 | 83 | 88 | - | - | 0 | Left frontal, parietal, medial |
| UMCU21 | 12 | F | 39 | 51 | 56 | 0 | 6 | 6 | Right frontal, parietal, occipital, medial |
| UMCU22 | 9 | F | 67 | 78 | 80 | - | - | 0 | Left frontal, parietal, temporal, medial |
| UMCU23 | 14 | F | 45 | 53 | 56 | - | - | 0 | Right frontal, parietal, medial |
| UMCU25 | 16 | M | 41 | 75 | 80 | 0 | 6 | 6 | Right frontal, parietal, occipital, temporal |
| UMCU26 | 7 | M | 45 | 57 | 64 | - | - | 0 | Right frontal, medial |
| UMCU59 | 50 | F | 41 | 47 | 48 | 14 | 16 | 16 | Right frontal, temporal |
| UMCU62 | 8 | M | 64 | 74 | 80 | - | - | 0 | Left frontal, parietal, temporal, medial |
| UMCU67 | 6 | F | 66 | 80 | 80 | - | - | 0 | Right frontal, parietal, temporal |

### 4.2 Electroencephalography recordings and stimulation

Single pulse electrical stimulation was performed with various subdural electrode grids and strips placed on the cortex of each of the 14 patients, as well as additional depth electrodes in six of the patients (Ad-Tech, PMT and DIXI). Subdural ECoG measurements at both Mayo Clinic and the UMC Utrecht were performed using electrode arrays with circular platinum contacts (2.3 mm exposed diameter, 10 mm center-to-center inter-electrode distance) embedded in a silastic sheet. Three patients at Mayo Clinic and three patients at the UMC Utrecht also had depth electrodes, where each lead provided 4, 6 or 8 cylindrical contacts at a 5mm inter-electrode distance. At both sites, signal data was amplified and sampled at 2048 Hz. Data at Mayo Clinic was recorded using a Natus Quantum amplifier and filtered between 0.01 and 878Hz. Data at the UMC Utrecht was recorded using a MicroMed LTM64/128 express electroencephalography headbox with integrated programmable stimulator.

Single pulse stimulation at Mayo Clinic was performed with a Nicolet cortical stimulator using a single biphasic 200 microsecond pulse on pairs of adjacent electrodes with a current of 6 mA. Pulses were triggered manually at an interval of ∼4 seconds. At the UMC Utrecht, single pulse stimulation used a monophasic 1000 microsecond pulse on pairs of adjacent electrodes with a current of 4 mA and 8 mA, at a 5 second inter-stimulus-interval.

Electrodes were excluded from analysis if they were not positioned on brain tissue in the case of ECoG (e.g. when one grid is partially on top of another grid); or not inside the brain in the case of SEEG. Electrodes were also excluded if, upon visual inspection, they were found to contain severe noise. Epochs with artifacts were marked in the data and excluded from analyses. Electrode-pairs were only included in analyses if they were stimulated 5 or more times (and thus included 5 or more trials).

The validation of the CCEPs was focused on ECoG data, given that these data have currently been shared in large databases ^60^. Section 2.3 shows an example that focuses specifically on SEEG, in which positive deflections are detected alongside negative deflections.

### 4.3 BIDS curation

All data were curated according to the BIDS iEEG specification. Following the example in the BIDS specification, each ‘_events.tsv’ file contained a column called ‘trial_type’ with the value ‘electrical_stimulation’ for each single pulse. Each ‘_events.tsv’ file also contained a column called ‘electrical_stimulation_site’ to describe the electrode pair that was stimulated (e.g. ‘CH01-CH02’). These ‘trial_type’ and ‘electrical_stimulation_site’ columns are required for the workflow to function correctly.

### 4.4 Time-locking, epochs, and normalization

We consider a CCEP response to be the response measured in one electrode after one other pair of electrodes is being stimulated (one trial). An average CCEP is the CCEP measured in one electrode averaged across multiple trials (median: 10; less than 5 were excluded) when another pair of electrodes is stimulated. To calculate average CCEPs, trials were time-locked to the stimulation onset and epoched/segmented with a (default) time-window of 1s before, to 2s after, stimulus onset. Each trial was normalized separately to the median of a baseline period of -1000ms to -100ms (i.e. before stimulus onset). Both the manual annotation and the automatic detection were performed on CCEPs for sets of trials that belonged to a single measured electrode given electrical stimulations on a specific electrode-pair. The interval that was affected by the stimulation artifact (0 to 9ms) was excluded from investigation. For each study this interval should be determined according to the hardware and can be configured in ER-detect.

### 4.5 Manual annotation of negative evoked responses

Manual annotation of 14 datasets (total of 59178 averaged responses) was performed by visual inspection of the average responses, and based on the presence of a negative deflection in the average response in each of the measured electrodes for each of the individual stimulated electrode-pairs. A negative evoked response was defined as a typical peak occurring within the interval between 10 and 90ms after stimulation onset. Before annotating the datasets at the Mayo Clinic, guidelines were established, and reviewed by clinical neurologists (NG, BL); The guide was added to this paper as Supplementary Table 2.

The data from the University Medical Center Utrecht were annotated independently by two researchers, while the data from the Mayo Clinic were annotated separately by another two researchers. Some sets were annotated double (intra- and inter-site) to be able to calculate inter-rater reliabilities. Table 4 provides an overview of the datasets, sites and the individual researchers that annotated the data.

**Table 4.**
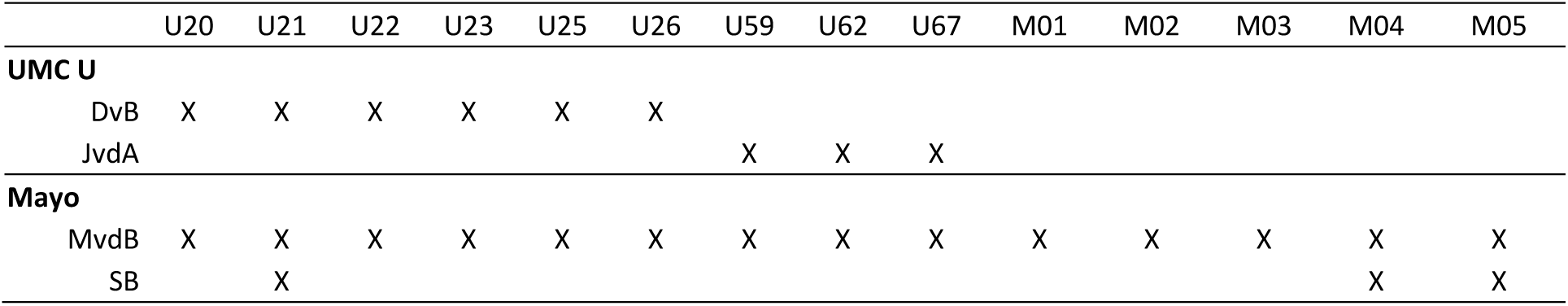
The visual inspection and manual annotation of datasets by researchers from both sites.

Two of the datasets were not completely annotated. One of the 41 stimulated electrode-pairs for subject UMCU21 was not annotated by rater DvB; and 14 of the 56 stimulated pairs for subject UMCU59 were not annotated by rater JvdA.

Although visual inspection is still a subjective interpretation of signal characteristics, having the annotations from four people from two independent sites allowed us to calculate the inter-rater reliability, which gives an indication of a rater-ceiling. Having a rater-ceiling is important for the comparison between the manual-visual inspection and automatic detection, as it provides somewhat of a maximum for automatic detection.

### 4.6 Automatic detection methods

Currently, three detection methods are implemented for the automated detection of evoked responses, based on (1) deviations from baseline, (2) inter-trial similarity, and (3) a wavelet filter. Detection is performed on each unique combination of a measured electrode and stimulated electrode-pair. Automatic detection always starts by effectively finding the negative peak amplitude (i.e. local extrema) in the average signal over trials, within a specific timespan after stimulation (9 to 90ms by default). If indeed a peak is found, one of the three methods can be used to determine whether that peak is classified as an evoked response.

By default, ER-detect uses the method that compares the peak amplitude to the standard deviation in a baseline period (Method 1 described in detail below). This method was chosen as a default because it is similar to what most previous studies implemented for automatic early response detection (see Table 1; ^2,5,59,60,64,15,34,47–49,52,55,58^). The other two methods (inter-trial similarity or the wavelet filter) can either be used for automated evoked response detection instead of, or in addition to the deviation from baseline method.

#### Method 1: Deviation from baseline

This method calculates the average evoked response across trials. From this average response, it then calculates the standard deviation during a baseline interval (default from -1000 to -100 ms). Similar as in other work ^47,60^, it is then determined whether the local extremum in the average signal exceeds a given amplitude threshold in order of standard deviations from baseline. The default threshold is set to 3.4 times the standard deviation but can be adjusted using a JSON file with settings. Similarly, the algorithm can be set to detect both negative and positive peaks.

#### Method 2: Inter-trial similarity

While the deviation from baseline method only depends on the average CCEP, an average CCEP can potentially be driven by a few outlier trials. We therefore added an optional method that is based on inter-trial similarity. Cross-projection of the trials is used to determine the inter-trial similarity (based on the Canonical Response Parameterization methods described by Miller et al., 2023 ^71^). In brief, the CCEPs measured in one electrode after stimulating one electrode pair *K* times can be represented in a matrix *V* with dimensions *T*×*K*, with *T* the total number of timepoints and *K* the total number of trials. A cross projection matrix *P*, is then calculated as scalar projections between all pairs of trials from the N1 interval (e.g. 12-90 ms), with self-projections removed: *P* = ^*V*^*T*^*V*, where 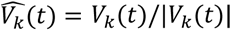 is unit normalized for each trial. The full matrix *P* is sorted into a combined set *S* with self-projection omitted, providing a distribution of cross-projections magnitudes across trials, which can be thought of as a measure for mutual information between trials. Then, it can be tested whether *S* is significantly larger than zero using a right tailed t-test. The resulting t-value(s) can be included as an additional output metric, or can be used as the primary method to automatically detect evoked responses. When this method is used to identify evoked responses, the resulting t-value is compared to a specific threshold. If it exceeds the threshold, then the deflections in the trials are considered similar enough and the peak is therefore identified as an evoked response.

#### Method 3: Wavelet filter

A typical early response in the average CCEP has an S-shape, which we often look for when performing visual inspection. This S-shape has a period of about 50 ms, and therefore has high power around 20 Hz. A wavelet filter is performed on the average signal of stimulus trials for each measured electrode given a specific electrode-pair. The average signal between 12-90ms after stimulus onset is isolated, re-centered by subtracting the mean over time and bandpass filtered from 10 to 30Hz using a third-order digital Butterworth filter. After filtering, the power is calculated using a Hilbert transformation.

Similar to the inter-trial similarity analysis, the resulting power value(s) can be included in the output as an additional metric or can be used as the primary method to automatically detect evoked responses. If the wavelet method is used to identify evoked responses, the resulting squared power value is compared to a specific threshold; when it exceeds the threshold, then the S-shape is pronounced enough to identify the peak deflection as an evoked response.

### 4.7 Inter-rater reliability and the validation of automated detection

The reliability of the manual annotations between multiple raters and validity of the automated detection methods were quantified in the same manner, by comparison of matrices. Both manual annotation and automatic detection provide a matrix that contains - for each measured electrode and each stimulated electrode-pair - whether an evoked response was found or not. To establish inter-rater reliabilities, the annotation matrices of different researchers on the same dataset were compared. To validate each of the automated detection methods, the annotation matrix of a researcher was compared to an output matrix from the tool. Annotations or detected responses that were found on measured electrodes within a 12mm radius of stimulated electrodes were excluded before comparison to rule out conduction through the cerebral spinal fluid (CSF). Upon comparing matrices, only the values of measured and stimulated electrodes that occur in both matrices were taken into account. The resulting sensitivity, specificity and Cohen’s Kappa values were used as indicators of the inter-rater reliability and validate the detection method.

## Data Availability

The data from Mayo Clinic and UMC Utrecht that was used in this article to validate the early evoked response (i.e. N1s) detection has been de-identified and is shared at: https://openneuro.org/datasets/ds004774

## Code Availability

The source code that is used to validate and visualize the data in this article is shared at: https://github.com/MaxvandenBoom/Paper_VandenBoom_ERDetect

The ER-Detect tool is shared at:

https://github.com/MultimodalNeuroimagingLab/erdetect (source-code, open under GPL 3.0)

https://pypi.org/project/erdetect/ (public python package)

https://github.com/MultimodalNeuroimagingLab/erdetect/wiki (documentation)

The ieegPrep library (part of ER-Detect) is shared at:

https://github.com/MultimodalNeuroimagingLab/ieegprep (source-code, open under GPL 3.0)

https://pypi.org/project/ieegprep/ (public python package)

https://github.com/MultimodalNeuroimagingLab/ieegprep/wiki (documentation)

## Acknowledgements

We thank Jaap van der Aar and Sam Buchl for their help in curating the data. For their contributions and help in collecting the data, we thank Cindy Nelson and Karla Crocket at the Mayo Clinic and the SEIN-UMCU RESPect database group at the UMC Utrecht (C.J.J. van Asch, L. van de Berg, S. Blok, M.D. Bourez, K.P.J. Braun, J.W. Dankbaar, C.H. Ferrier, T.A. Gebbink, P.H. Gosselaar, M.G.G. Hobbelink, F.W.A. Hoefnagels, N.E.C. van Klink, M.A. van ‘t Klooster, G.A.P. de Kort, M.H.M. Mantione, A. Muhlebner, J.M. Ophorst, R. van Regteren, P.C. van Rijen, S.M.A. van der Salm, E.V. Schaft, M.M.J. van Schooneveld, H. Smeding, D. Sun, A. Velders, M.J.E. van Zandvoort, G.J.M. Zijlmans, E. Zuidhoek and J. Zwemmer).

## Author Contributions

Max van den Boom - Study design, data curation/analysis/interpretation, development of software, writing and revising the manuscript.

Nick Gregg - Data acquisition and interpretation and revising the manuscript.

Gabriela Ojeda Valencia - Data acquisition, curation, annotation and revising the manuscript.

Brian Lundstrom - Study design, data interpretation and revising the manuscript.

Kai Miller - Data acquisition and revising the manuscript.

Dorien van Blooijs - Data acquisition, curation, annotation and revising the manuscript.

Geertjan Huiskamp - Data acquisition and revising the manuscript.

Frans Leijten - Study design, data acquisition and revising the manuscript.

Greg Worrell - Study design, data acquisition and revising the manuscript.

Dora Hermes - Study design, data acquisition, analysis, interpretation, writing and revising the manuscript.

## Competing Interests

Research reported in this publication was supported by the National Institute of Mental Health of the National Institutes of Health under Award Number R01MH122258 (DH, MvdB, GOV, FSSL, GAW, KJM; the content is solely the responsibility of the authors and does not necessarily represent the official views of the National Institutes of Health), the Mayo Clinic DERIVE Office and Center for Biomedical Discovery support (DH, KJM, MvdB, GAW), and the Epilepsy Foundation of the Netherlands under Award Number NEF17-07 (DvB).

BNL has no personal financial interests, but declares intellectual property licensed to Cadence Neuroscience Inc (contractual rights waived; all funds to Mayo Clinic) and Seer Medical Inc (contractual rights waived; all funds to Mayo Clinic), site investigator (Medtronic EPAS, Neuroelectrics tDCS for Epilepsy), industry consultant (Epiminder, Medtronic, Neuropace, Philips Neuro; all funds to Mayo Clinic), and educational support (Dixi Medical). NMG declares industry consultant for NeuroOne Inc., funds to Mayo Clinic. GAW was supported by National Institutes of Health Grant UH2&3 NS095495 and unrelated to this research has licensed intellectual property developed at Mayo Clinic to Cadence Neuroscience Inc. and NeuroOne Inc.

## Notes

https://openneuro.org/datasets/ds004774

