## Supplementary Table 1 & 2 and Supplementary Figure 1 for "ER-detect: a pipeline for robust detection of early evoked responses in BIDS-iEEG electrical stimulation data"

### Supplement Table of Contents

- Supplementary Table 1. Literature based definitions of N1 and other evoked components

- Supplementary Table 2. Guide used to annotate N1s

- Supplementary Figure 1. ROC curves per subject

#### Supplementary Table 1. Literature based definitions of N1 and other evoked components

In this manuscript, we present a tool to detect evoked responses in iEEG data. When measured on the surface with ECoG electrodes, these responses often show a negative deflection between 9 and 50 ms, which has been described as an N1 response in many studies. We validate the ER-detect tool detection of negative evoked (N1) responses and compare this to visual annotations.  This table reviews how previous literature has described N1 responses. In the next supplement, we describe a guide for how we visually annotated the responses in our data for validation. Note that when annotating our data, we also observe other response shapes. These include later responses and early positive responses. While the tool can be used to detect any early responses whether positive or negative, we only annotated and validated N1 responses.

**P*:** In the literature:
- Some authors (Conner, 2011) label P1 as the change that occurs after the N1.
- The Cleveland group (Nair, Matsumoto)  (Araki, 2015; Terada, 2012; Matsumoto 2017) name the P1 as "the 'very first volley' that occurs before the N1 (which in practice is often masked by the stimulation artifact)." They also acknowledge the second positive deflection but use variable labels or no label.

All images in Supplemental Table 1 were drawn to schematically represent the responses published in each article, and do not contain any source images.

| Article | Images | N1 (& N2) in text | Positive deflection(s) in text |
|---|---|---|---|
| Almashaikhi et al. (2014) [31] | 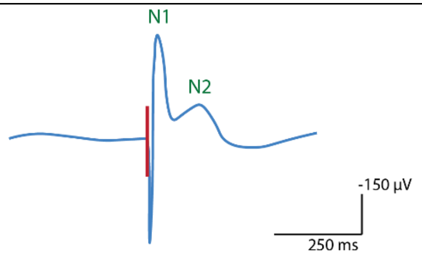  Reconstruction of Fig 1 | • "curves are the average of two 20 trials, showing similar N1 and N2 peaks" | n/a |
| Araki et al. (2015) [9] | 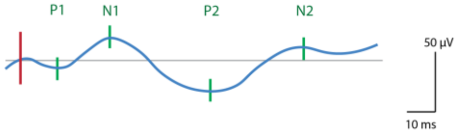  Reconstruction of single trial from Fig 1 (LBP4) | • "According to a previous report (Matsumoto et al., 2004), CCEPs consist of an early (N1) and a late (N2) negative potential. We measured the latency and voltage of N1 and N2." | • "The positive potential preceding N1 was designated P1, and that following N1 was named P2." |

| | | | |
|---|---|---|---|
| Conner et al. (2011) [10] | 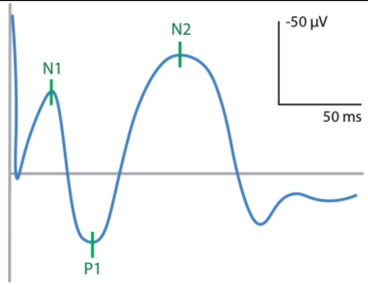<br>Reconstruction of Fig 2C | • "The first clear negative deflection following the stimulus artifact was defined as an N1 response [Matsumoto et al, 2004, 2007] (Fig. 2)."<br>• "For the vast majority of electrodes analyzed in this study, the N1 potential was the first and only potential identified."<br>• "... the N1 evoked potential. Slower or subsequent peaks in the evoked potential are not considered." | n/a |
| Enatsu et al. (2012a/2012b/ 2013) [1,43,44] | 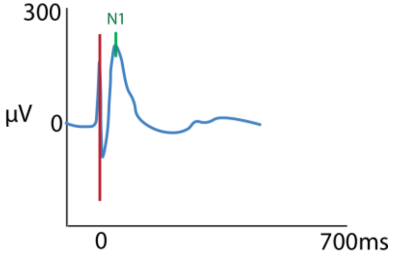<br>Reconstruction of Fig 2A<br>(iCCEP; Enatsu, 2012, Clin. Neuro.) | • "The N1 peak was visually identified as the first negative deflection that was clearly distinguishable from the stimulus artifact." (2012a/2012b/2013a) | n/a |
| Entz et al. (2014) [52] | 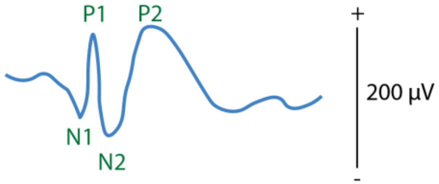<br>Reconstruction of Fig 2 | • "N1: 10–50 ms, N2: 50–500 ms" | n/a |
| Greenlee et al. (2004) [53] | 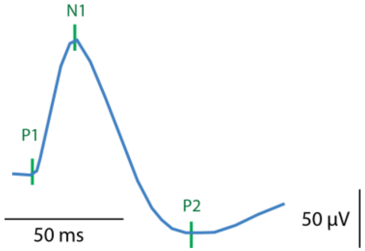<br>Reconstruction of Fig2B (20V stim. trace) | • "Major deflections are referred to as P1, N1, and P2 to indicate deflection polarity in order of appearance." | |

| | | | |
|---|---|---|---|
| Howard et al. (2000) [26] | 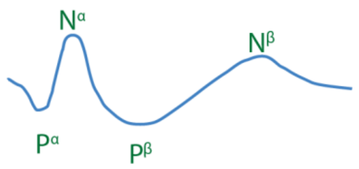<br>Reconstruction of Fig 2 | • "The EP was characterized by an initial positive peak (Pa) followed by a negative component (Na) and then by additional positive (Pb) and negative waves (Nb) of smaller amplitude." | |
| Kamada et al. (2020) [54] | n/a | • "We paid the most attention to assign and confirm consistent CCEP findings. CCEP waveforms generally demonstrate 2 negative peaks (N1 and N2)..." | n/a |
| Kanno et al. (2018) [13] | 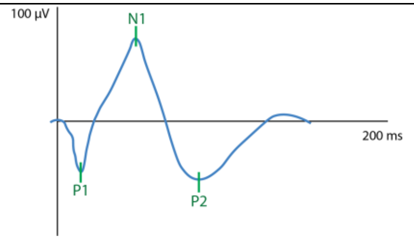<br>Reconstruction of Fig1B | • "Typical CCEPs consist of an early (N1) and a late (N2) negative potential (Matsumoto, 2004)."<br>• "N1 peak was visually identified as a negative deflection that was clearly distinguishable from the stimulus artifact." | n/a |
| Keller et al. (2011, 2014) [2,55] | 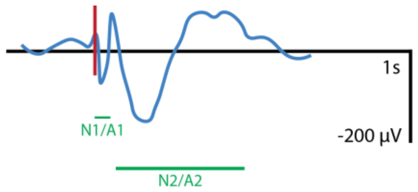<br>Reconstruction of Fig1A (2011) and Fig1A (2014) | • "The early CCEP consists of two major deflections (usually negative in polarity) termed N1 and N2, peaking at ~20–30 ms and 120 ms, respectively." (2011)<br>• "CCEPs in human cortex generally consist of an early sharp response (10–50 ms post stimulation) and a later slow-wave (50–250 ms). These responses have previously been referred to as N1 and N2, respectively… (Matsumoto et al., 2004). " (2014) | • "For comparison, we also computed CCEPs for the P1 (20–50 ms) and N1 (20–50 ms) time windows…" (2011) |
| Lemarechal et al. (2022) [48] | n/a | • "CCEPs generally consist of a first sharp peak (10–50ms), the N1 component, followed by a slow wave (80–250ms), the N2 component."<br>• "only significant CCEPs with a peak latency comprised in the first 80ms were selected, in order to limit the analysis to the early N1 component" | • "this N1/N2 terminology is a simplification of the complexity of CCEPs, which also includes positive components" |

| | | | |
|---|---|---|---|
| Matsumoto et al. (2004, 2007, 2012) [3,16,56]<br><br>Koubeissi et al. (2012) [14]<br><br>Ookawa et al (2017) [17] | 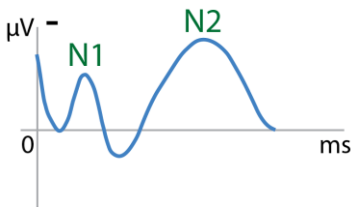<br>Reconstruction of Fig 1 (Matsumoto et al, 2004) | • "The N1 peak was visually identified as a first negative deflection that was clearly distinguishable from the stimulus artefact." (Matsumoto et al., 2004, 2007, 2012; Ookawa et al, 2017)<br>• "We measured the voltage of the CCEPs in a manner identical to Matsumoto et al. (2004)" (Koubeissi al., 2012)<br>• "The onset, peak latency, and amplitude of N1 were measured as previously reported. Matsumoto et al., 2004)" (Ookawa et al, 2017) | n/a |
| Matsumoto et al. (2017) [57] | 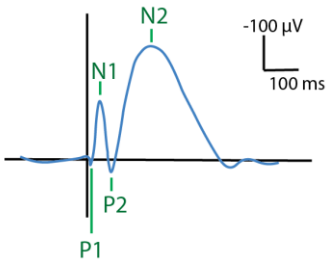<br>Reconstruction of Fig4C.<br>Same as Usami et la. (2015). | • "Initial studies found that CCEP generally consists of an early sharp negative potential (N1: peak 10–50 ms) and a later slow- wave like potential (N2: peak 50–300 ms) [Matsumoto et al, 2004, 2007]"<br>• "N1 could be recorded as a positive potential, potentially reflecting the positive end of the dipolar activity in the sulcus [Matsumoto et al., 2004; Keller et al.]."<br>• "In some case, N1 peak latency could exceed 50 ms [Araki, 2015], and the N2 potential (peak latency of >100 ms) may occur alone." | • "The troughs or positive sharp reflections before and after the N1 peak were termed P1 and P2, respectively"<br>• "A small positive deflection preceding N1, namely, the onset of N1, is termed P1 by some authors since this could reflect the very first volley to the target cortex [Terada et al, 2012, 2008]." |
| Nakae et al. (2020) [15] | 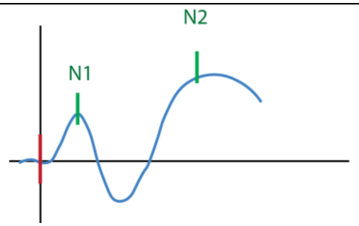<br>Reconstruction of Fig3 (R1 aMTG/aITG x IFG pOrb) | • "…the N1 cluster determined in this method was similar to the traditional criteria of N1 (onset < 30 ms, peak < 100 ms)." | n/a |
| Silverstein et al. (2020) [58] | 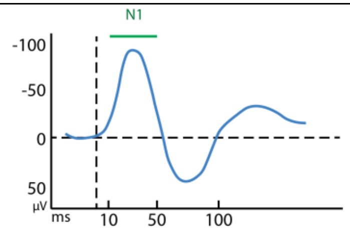<br>Reconstruction of Fig 2. | • "The N1 amplitude and latency were defined as the peak negative voltage within the 10–50 ms post-stimulus window" | n/a |

| | | | |
|---|---|---|---|
| Suzuki et al. (2019) [19] | 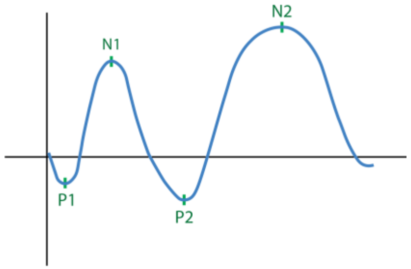  Reconstruction of Fig 1. | • "The N1 peak was visually identified as a first negative deflection that was clearly distinguishable from the stimulus artefact." (same as Matsumoto et al., 2004) | • The initial positive peak (P1) preceded N1 responses in 2 patients. N2 response was not observed in 4 patients. |
| Terada et al. (2012) [8] | See Fig 2-4, 5 and 6 in the article. | • "As reported previously, initial positive triphasic waves (P1-N1-P2) were recorded from the contralateral hemisphere (Figs. 2–4; Type 1 response)."<br>• "...initial negative biphasic waves (N1-P2) were observed (Fig. 5; Type 2 response)."<br>• "In addition, initial positive biphasic waveforms (P1-N1) were identified in 8 recordings (Fig. 6; Type 3 response)." | |
| Trebaul et al. (2018) [49] | 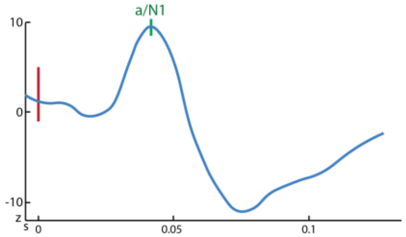  Reconstruction of Fig 1. | • "..two kinds of approaches are currently used to identify the response components.": 1)"... from visual analysis and described as two peaks (N1 and N2) ..." or 2) "More recently, statistical tests have been proposed...". " ... baseline normalization and the use of a threshold on the z-score were chosen here."<br>• Statistical first peak (latency = a/N1) corresponds to visual N1 | n/a |
| Umeoka et al. (2009) [21] | 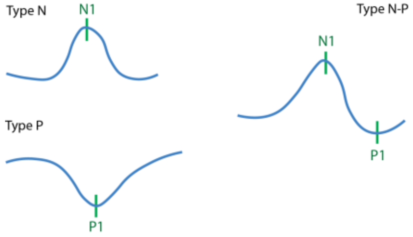  Reconstruction of Fig 2 | • "We classified the CCEP waveforms into 3 types: type N- P ... consisted of an initial negative peak (designated N1) followed by a positive peak (designated P1), type N consisted of N1 without P1, and type P consisted of P1 only (Fig. 2)." | |

| | | | |
|---|---|---|---|
| Usami et al. (2015) [59] | 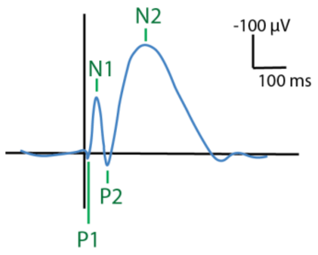<br><br>Reconstruction of Fig 1D.<br>Same as Matsumoto et al. (2017). | ● "CCEP often comprises first sharp (N1) and then slow (N2) negative components as previously mentioned. The troughs or positive sharp reflections before and after the N1 peak were termed P1 and P2, respectively (Fig.1D)."<br>● "N1 was clearly distinct from both the stimulus artifact and N2" | |
| Van Blooijs et al. (2018) [47] | n/a | ● "Within 100 ms after the stimulus, early responses (ERs) may be observed … this ER is known as the N1-response (Enatsu, Piao, et al., 2012; Keller et al., 2014; Matsumoto et al., 2004, .2007)" | n/a |
| Vega-Zelaya et al. (2023) [22] | 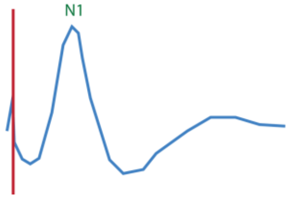<br><br>Reconstruction of Fig 1C. | ● "A positive response was recorded if a large N1 peak (upward directed) in the electrodes of the PLA was obtained (Matsumoto, 2004; Yamao, 2014)" | n/a |
| Yamao et al. (2014, 2017) [23,24] | n/a | ● "The N1 peak was visually identified as the first negative deflection that was clearly distinguishable from the stimulus artifacts." (2014)<br>● " The onset, peak latency, and amplitude of N1 were measured as reported previously … [Matsumoto et al., 2004]." (2014, 2017) | n/a |
| Zhao et al. (2019) [61] | n/a | "Each CCEP consists of an early sharp negative response (N1, 10–50 ms post-stimulation) and a subsequent slow-wave (N2, 50– 300 ms post-stimulation) (Matsumoto et al., 2017). Here we only focused on the earliest response." | n/a |

#### Supplementary Table 2. Guide used to annotate N1s

Please note that negative is downward in this supplement

| Characteristics | Example(s) | Annotate | Explanation |
|---|---|---|---|
| | **Clear responses** | | |
| Clear negative deflection | 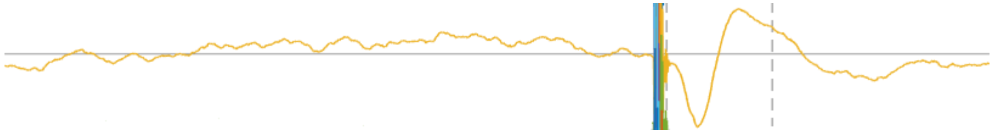 | N1 | - |
| Only positive deflection | 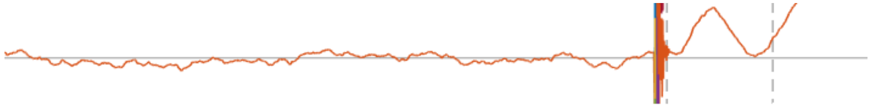 | No N1 | We only validate early negative responses, as these positive responses are not extensively documented in previous literature |
| Positive deflection followed by a negative deflection | 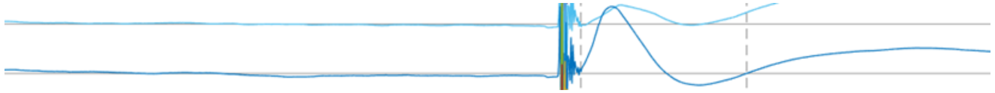 | No N1 | The latter negative deflection cannot be seen as the initial response. |
| | **Noisy responses (stimulation artifacts and drifts)** | | |
| Strong early response, often next to stim electrode | 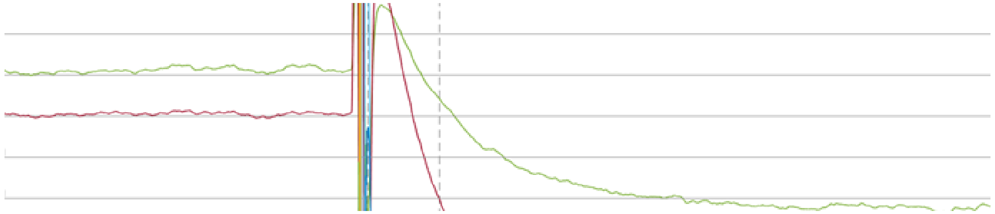 | If severe, do not annotate<br><br>If not severe, annotate | Most likely the result of stimulation leaking over from stim electrode |

| | | | |
|---|---|---|---|
| Shows left slope of ER, but keeps on drifting after peak | 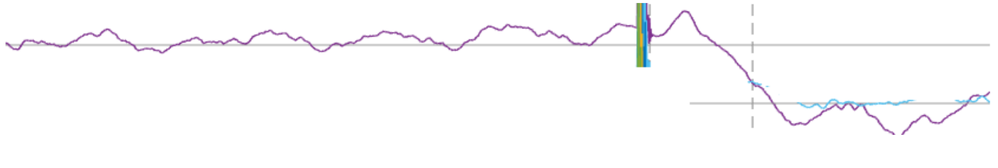 | No N1 | If the latter drifting part has no typical ER shape (very wide) or falls far outside of the time range (very late), then that drift is not an ER |
| Already an artifact before stim onset. | 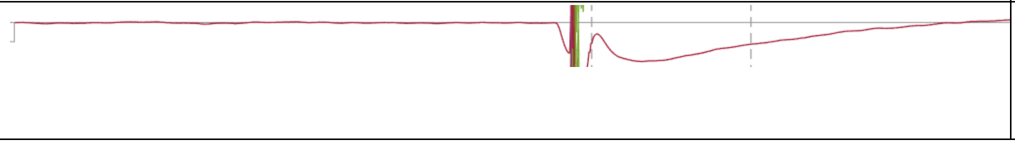 | Do not annot. | Too much artifact in signal (could be leakage/close to stim electrode) to judge whether it actually comes from the brain |
| **Complex responses (pushed by stim)** | | | |
| Stim pushes signal up/down, but no distinct ER shape after | 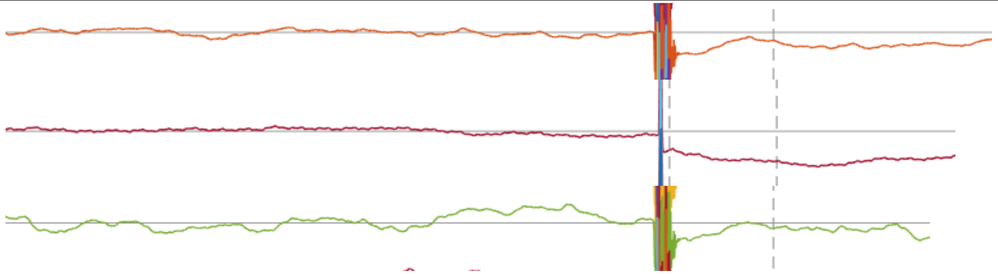 | No N1 | *Just shift of baseline due to stim [ref]* |
| N1 flowing from stim, can still see N1 going down in stim artifact | 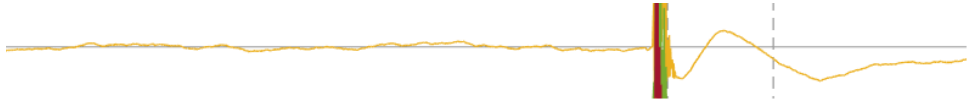 | N1 | *Typical N1, the response is so early it coincides partially with the stimulation artifact* |
| The N1 peak seems to be so early, during the stimulation artifact (<9ms) | 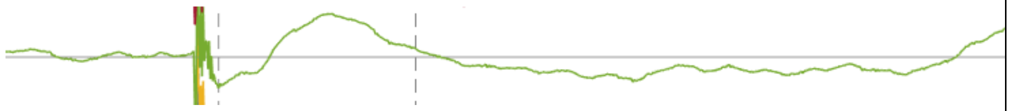 | No N1 | *Too early to measure, don't consider responses before the end of stimulation artifact (~9ms).* |
| Clear ER, pushes up to be an Inverted N1, but then drops enough to be considered a N1 as well. | 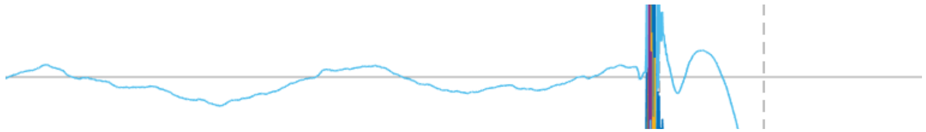 | N1 | Annot on either direction, even if typical N1/Class 2 is later (after push). Reference is still baseline before stim. |

| | | | |
|---|---|---|---|
| **Complex responses (fluctuating baseline)** | | | |
| Strong (+-500uV) fluctuations in signal (also before stim) with no clear response | 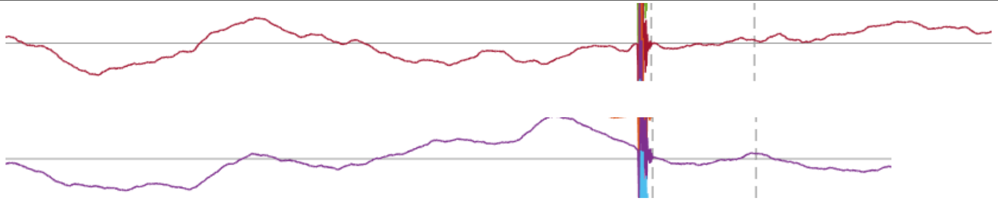 | No N1 | - |
| Strong/medium fluctuations in signal but distinct ER shape that differs from earlier fluctuations | 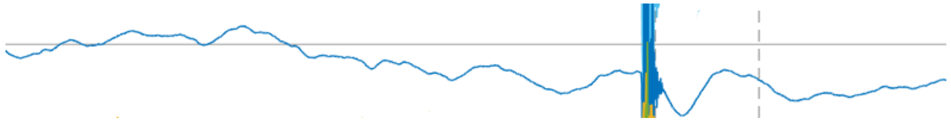 | N1 | *Noisy signal, but no reason not to annotate since a response does occur* |
| Signal already goes up/down before stim. | 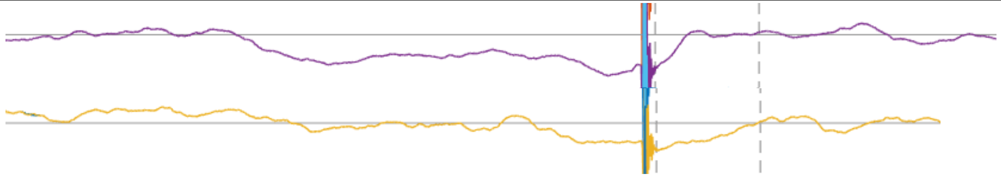 | No N1 | *Assume it is normal signal movement coinciding with stim* |
| **Complex response (timing, size or direction)** | | | |
| Smaller yet distinct N1 shape | 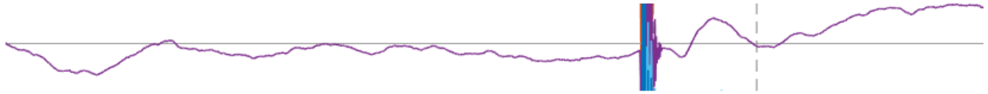 | N1 | - |

| | | | |
|---|---|---|---|
| Seemingly ER that occurs just after stim, but which shape also occurs before the stim. | 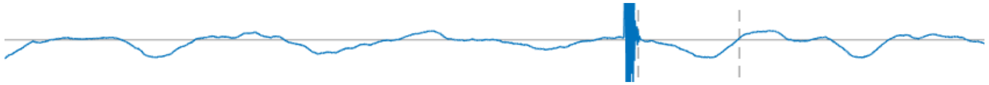 | No-N1 | Fluctuation same in baseline |
| Signal below zero | 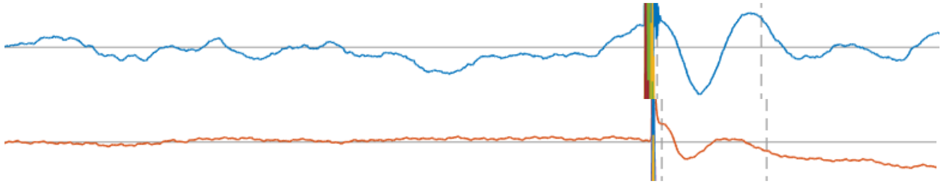 | N1 | *The baseline before stimulus remains the reference (0), elsewise the phasing will determine reference.* |
| Signal not below zero | 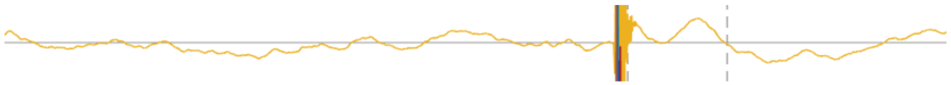 | No N1 | Annot, with baseline before stim as ref.<br><br>*The baseline before stimulus remains the reference (0), elsewise the phasing will determine reference.* |
| ER that is not within reasonable range or large width, but with typical ER shape | 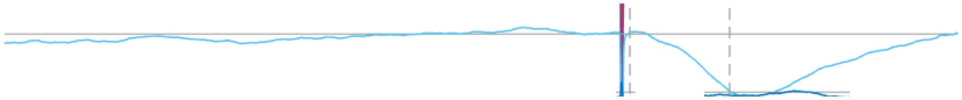 | No N1 | ER if spans < 80ms and is reasonably within time-range. A very wide (e.g. 200ms) N1 is No-ER<br><br>*Example: Too wide, No-ER* |

#### Supplementary Figure 1. ROC curves per subject

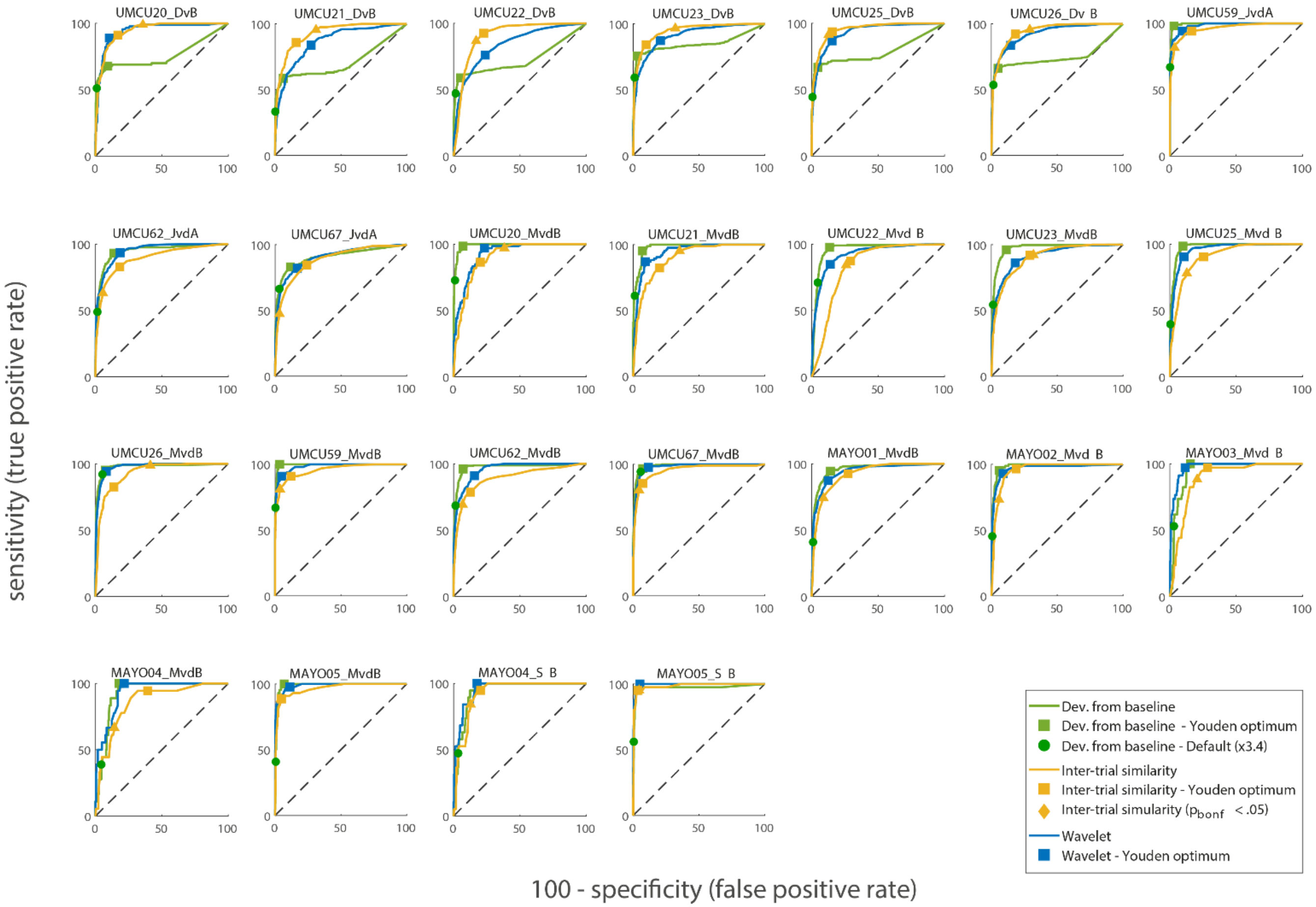


***Supplementary Figure 1. Performance of the three different detection methods per subject.*** *ROC curves projecting the true positive rate (sensitivity) and false positive rate (specificity) for each automated detection method. The 'deviation from baseline' method is shown in green with the threshold (as a factor of the standard deviation during baseline) ranging from 1 to 1500. The green dot represents this method's performance based on the default setting of 3.4. The 'inter-trial reliability' performance is shown in yellow with threshold t-values ranging from -30 to 30. The 'wavelet' performance is shown in blue with power thresholds ranging from 0 to 60 000.*
